# Quantitative AI-based DNA fiber workflow to study replication stress

**DOI:** 10.1101/2025.03.27.645743

**Authors:** Paolo Fagherazzi, Timo Diekmann, Alessandra Ardizzoia, Zuzana Machacova, Simran Negi, Anoop Kumar Yadav, Pavel Moudry, Vincenzo Costanzo, Hana Polasek-Sedlackova

## Abstract

Replication stress (RS) is a prominent source of genome instability and human diseases. Understanding its molecular mechanism through various quantitative and unbiased methodologies is essential for the advancement of treatment strategies. One of the powerful methods to study DNA replication dynamics and its alterations at the single-molecule resolution is the DNA fiber assay. However, this method relies exclusively on manual image acquisition and analysis, making it time-consuming and susceptible to user bias. Here, we present a quantitative AI-based DNA fiber (qAID) workflow enabling imaging and multiparameter analysis of thousands of DNA fibers within several dozen minutes. Our workflow quantifies key parameters, including DNA fiber frequency, length, and symmetry, while also allowing visual inspection of individual DNA fibers using unbiased image galleries. The robustness of the workflow is demonstrated by comprehensive datasets of biologically relevant experiments performed by three independent laboratories. Overall, qAID workflow provides a fast and effective examination of replication dynamics and its alterations at the single-molecule resolution.

## INTRODUCTION

The precise and complete duplication of genetic information during DNA replication is essential for each cell division. The initiation of eukaryotic DNA replication is divided into two main steps: origin licensing and firing, which occur in distinct phases of the cell cycle^1^. During origin licensing in the G1 phase, minichromosome maintenance 2-7 (MCM2-7) complexes are loaded onto replication origins as inactive double hexamers. In the S phase, a subset of the loaded MCM2-7 double hexamers is converted into two active CDC45-MCM2-7-GINS (CMG) helicases through the concerted action of specific CDK and DDK along with the firing factors^2^. The formation of CMG helicases provides a basis for the assembly of two replisomes, which contain core enzymes essential for DNA unwinding and synthesis of new strands. In addition to core enzymes, replisomes include various accessory components that coordinate enzymatic activities at the replication fork, ensuring accurate genome duplication.

During replication elongation, replication forks encounter a variety of challenges from both extrinsic and intrinsic factors that can disrupt the enzymatic activities of the replisome and eventually cause replication-associated DNA damage. Such perturbations at the replication forks are known as replication stress^3,4^. Depending on the level of RS, the cell can activate an adequate response mechanism. Mild RS, such as that caused by metabolic fluctuations, triggers transient and reversible replisome remodeling, leading to a temporary slowdown of replication fork progression^5,6^. Severe RS, induced by factors such as chronic dNTP depletion or chemotherapeutic agents, can lead to the bypass of DNA lesions through re-priming of DNA synthesis and the formation of daughter strand gaps (DSGs) or cause replication fork stalling^7-9^. The stalled replication forks can be actively reversed into a four-way junction DNA structure by specialized translocases, and upon RS release, reversed forks are restored, enabling the resumption of synthesis^10^. However, if replication forks become permanently stalled, this can result in their collapse, triggering break-induced replication repair and local activation of backup origins to rescue DNA replication^5^. In the absence of these protective mechanisms, DNA replication becomes a highly toxic and error-prone process, leading to DNA damage and genomic instability.

Extensive research over recent decades has elucidated RS as a major contributor to genomic instability and tumorigenesis^3,4^. Thus, a comprehensive understanding of the molecular mechanisms underlying RS, utilizing various quantitative and unbiased methodologies, is essential for the advancement of cancer treatment strategies. Quantitative image-based cytometry (QIBC), utilizing scanR high-content imaging system, has proven to be a highly effective technique for investigating RS at the single-cell level^11-15^. Among the various methodologies for analysis of DNA replication dynamics and its alterations at single-molecule resolution, the DNA fiber assay stands out^16-18^. This method allows for the detailed examination of critical parameters, including replication fork speed, symmetry of replication forks, and the frequency of different classes of DNA fibers^19^. However, unlike the quantitative assessment of RS at the single-cell level, the DNA fiber assay depends on manual acquisition and measurement of replication fork parameters. This reliance on manual techniques not only renders the approach time-consuming but also introduces the potential for user bias, which may compromise the accuracy and reproducibility of the findings.

In this manuscript, we present the development and comprehensive characterization of qAID workflow for the automated acquisition and analysis of DNA fibers using the scanR high-content imaging system. The workflow enables automated imaging, segmentation, and multiparameter analysis of thousands of DNA fibers within a short time frame. DNA fiber segmentation uses two neural networks (NNs) to classify fibers into five categories: origin firing, ongoing, stalled, and terminated fork. Following the classification, the multiparameter analysis extracts vital information, including the frequency, speed, and symmetry of individual replication forks. Additionally, visual inspection of individual DNA fibers is facilitated through unbiased image galleries, complementing the multiparameter analysis. In summary, the qAID workflow represents a fast and efficient method for examining replication dynamics and its alterations at a single-molecule resolution, enhancing our understanding of RS and related diseases.

## RESULTS

### Automated acquisition of DNA fibers using scanR high-content imaging system

The DNA fiber assay is one of the most powerful and widely used methods for studying DNA replication dynamics and its alterations at the single-molecule level^16-18^. This assay relies on the sequential labeling of newly synthesized DNA in living cells using two halogenated thymidine analogs, such as 5-chloro-2’-deoxyuridine (CldU) and 5-iodo-2’-deoxyuridine (IdU) (Fig. 1a). After sequential labeling, cells are harvested and subsequently processed by either DNA fiber spreading or combing (Fig. 1b). In case of DNA fiber spreading, the cells are subjected to *in situ* lysis to release DNA fibers on glass slides. The slide is inclined at an angle of approximately 15 degrees, allowing DNA fibers to stretch under gravity, leading to a non-uniform distribution across the slide^16,20^. In contrast, the DNA combing technique involves lysing labeled cells within agarose plugs, followed by stretching the DNA fibers on a silanized glass coverslip at a slow, constant rate with a combing machine^17,21^. This approach yields a parallel and uniform distribution of DNA fibers on the glass surface. After the DNA fibers are stretched using either method, they are fixed, denatured, and neutralized. The labeled DNA is visualized through immunofluorescence staining, utilizing antibodies that detect halogenated thymidine analogs incorporated into the DNA. For detailed DNA fiber spreading and combing protocol, please refer to the Method section of this manuscript.

**Figure 1:**
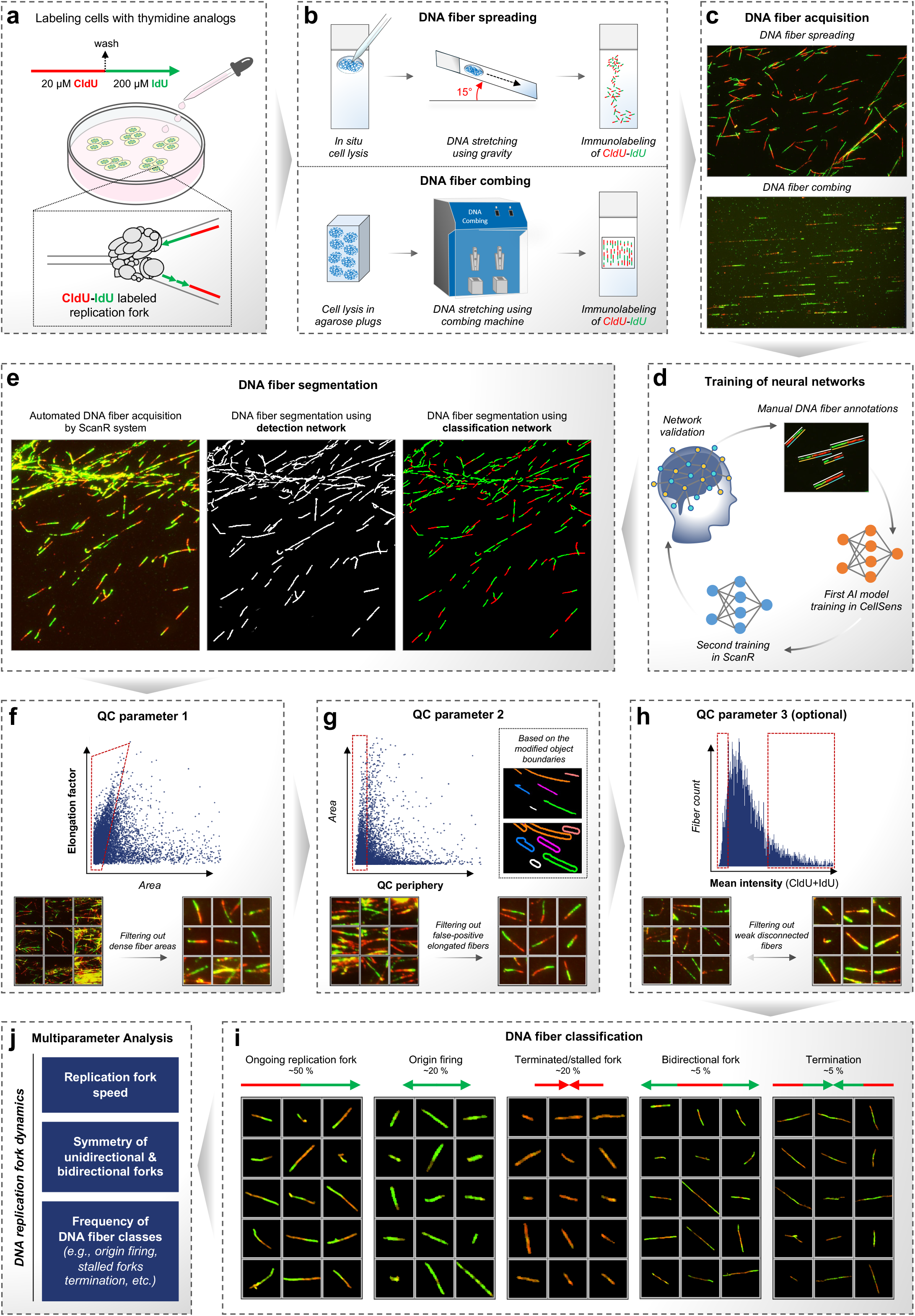
qAID concept and workflow. **(a)** A schematic representation illustrating the concept of the DNA fiber method. **(b)** A schematic representation illustrating the key steps in the preparation of DNA fibers by spreading and combing techniques. **(c)** Representative images of DNA fibers prepared by spreading and combing techniques. **(d)** A model summarizing the development of NNs for the detection and classification of DNA fibers (see text for details). **(e)** An example of DNA fiber segmentation using DNA fiber detection and classification NNs. The middle and right images are probability images created by the respective NNs. **(f)** Application of QC parameter 1 through plot displaying elongation factor and area parameters for individual DNA fibers. A red gate represents a gating strategy to filter out large, dense DNA fiber areas. The effect of the applied gating strategy is visualized by unbiased DNA fiber galleries. **(g)** Application of QC parameter 2 through plot displaying QC periphery and area parameters for individual DNA fibers. A red gate represents a gating strategy to filter out false-positively elongated DNA fibers. The effect of the applied gating strategy is visualized by unbiased DNA fiber galleries. **(h)** Application of QC parameter 3 through plot displaying the mean intensity of merged fluorescent channels and count parameters for individual DNA fibers. Red gates represent a gating strategy to filter out unfocused, weak, or disconnected DNA fibers. The effect of the applied gating strategy is visualized by unbiased DNA fiber galleries. **(i)** Classification of DNA fibers into five classes represented by unbiased qAID galleries. Galleries were masked based on segmentation. **(j)** A schematic representation illustrating the main features of multiparameter analysis to examine replication fork dynamics.

**Figure 2:**
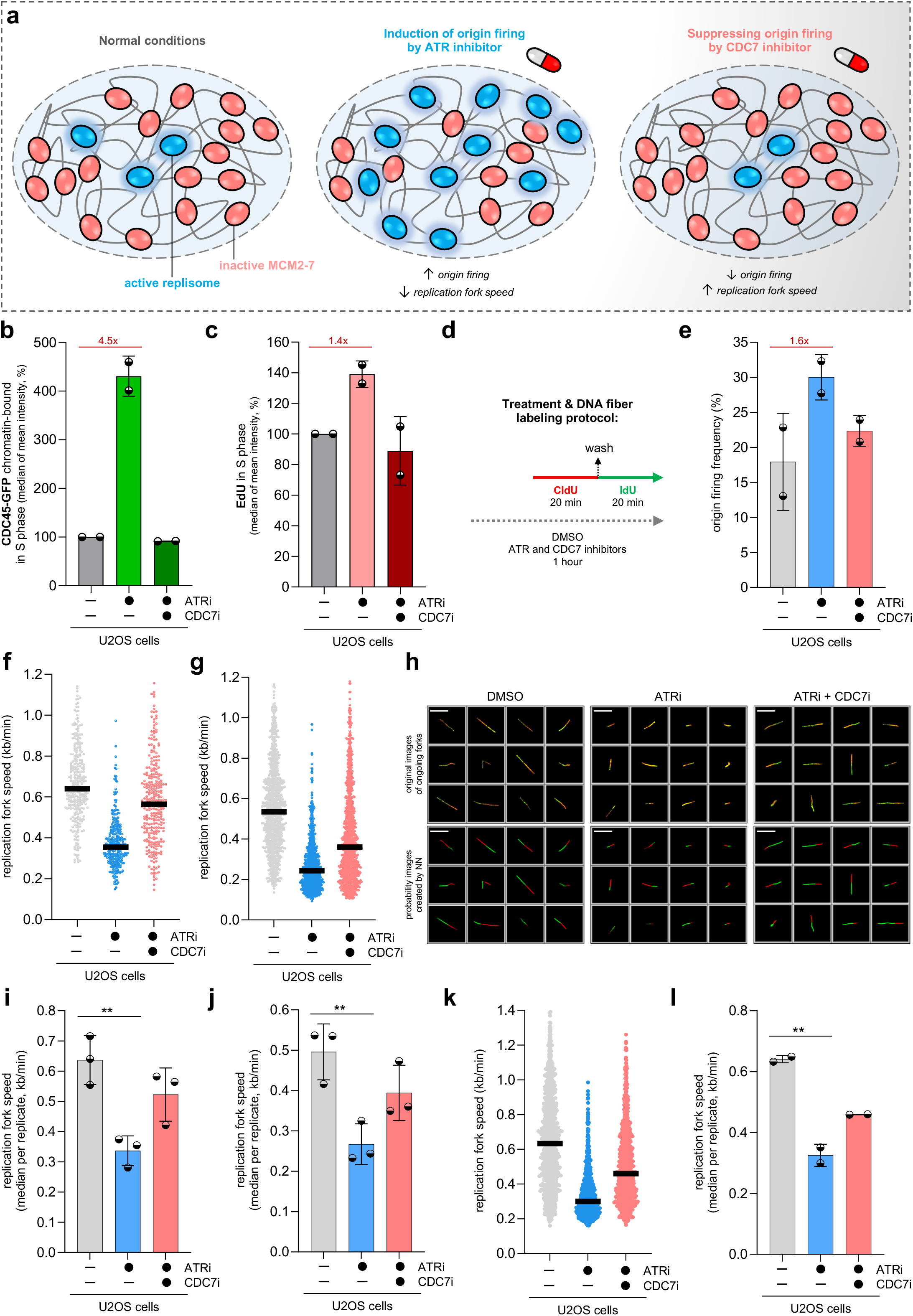
Quantitative analysis of the adaptation of replication fork speed in response to fluctuations of origin firing. **(a)** A schematic representation illustrating the concept of how the level of origin activation induced by ATR and CDC7 inhibitors affects the replication fork speed (see text for details). **(b)** Quantification of QIBC plots in Extended Data Fig. 3a; each bar indicates the median of mean intensity (data are mean ± s.d.; n = 2 independent experiments). **(c)** Quantification of QIBC plots in Extended Data Fig. 3d; each bar indicates the median of mean intensity (data are mean ± s.d.; n = 2 independent experiments). **(d)** DNA fiber labeling protocol with indicated DMSO, ATR, and CDC7 inhibitors treatment (5 µM ATRi, 5 µM CDC7i for 1 h). **(e)** qAID-based quantification of origin firing frequency in U2OS cells treated with indicated inhibitors (data are mean ± s.d.; n = 2 independent experiments). **(f)** Manual quantification of replication fork speed for ongoing forks in U2OS cells treated with indicated inhibitors. DNA fibers were prepared by spreading technique. Lines represent medians; n ≈ 350 fibers per condition. **(g)** qAID-based quantification of replication fork speed for ongoing forks in U2OS cells treated with indicated inhibitors. DNA fibers were prepared by spreading technique. Lines represent medians; n ≈ 1000 fibers per condition. **(h)** Unbiased qAID galleries of DNA fibers in **(g)**. Both original fiber images and probability images created by the classification network are displayed. Galleries were masked based on segmentation. Scale bar, 10 μm. **(i)** Quantification of replication fork speed plots in **(f)**; data are mean ± s.d.; n = 3 independent experiments. P value was calculated by a two-tailed unpaired *t*-test (P = 0.0053). **(j)** Quantification of qAID-based replication fork speed plots in **(g)**; data are mean ± s.d.; n = 3 independent experiments. P value was calculated by a two-tailed unpaired *t*-test (P = 0.0098). **(k)** qAID-based quantification of replication fork speed for ongoing forks in U2OS cells treated with indicated inhibitors. DNA fibers were prepared by combing technique. Lines represent medians; n ≈ 1000 fibers per condition. **(l)** Quantification of qAID-based replication fork speed plots in **(k)**; data are mean ± s.d.; n = 2 independent experiments. P value was calculated by a two-tailed unpaired *t*-test (P = 0.0073).

DNA fiber images can be acquired on any microscopy system (Fig. 1c), as demonstrated by our analysis of papers utilizing DNA fibers to study replication stress (Extended Data Fig. 1a, b). However, the manual acquisition of DNA fibers is time-consuming and yields relatively low throughput. In the past, the company Genomic Vision provided high-throughput services, such as Easy Scan, for scanning entire coverslips containing DNA fibers^21^. In this study, we utilized a scanR high-content imaging system equipped with infrared laser hardware autofocus to acquire DNA fiber samples. This setup facilitates precise and automated scanning of DNA fibers across entire coverslips or within user-defined regions, capturing thousands of DNA fibers in a matter of minutes. For further details on DNA fiber acquisition using the scanR high-content imaging system, please refer to the Method section or Supplementary File 1.

### Automated and quantitative AI-based analysis of DNA fibers

Although DNA fiber assay is a powerful and widely used method to study DNA replication dynamics at the single molecule resolution, it relies on manual image analysis (Extended Data Fig. 1c), which can be both time-consuming and prone to user bias. This bias can be manifested in several ways, such as inconsistencies in the segmentation of disconnected or spotty fibers (Extended Data Fig. 1d, e), unintentional omission of short DNA fibers (Extended Data Fig. 1f, g; 2a), or challenges in classifying asymmetrical DNA fibers (Extended Data Fig. 2b-d). Such an inter-user variability in manual quantification underscores the necessity for reliable computational methods that facilitate rapid and unbiased analysis of DNA fibers. To address this, we employed the TruAI deep learning module in CellSens and scanR software to train two distinct NNs. The first neural network, the DNA fiber detection NN dedicated to whole fiber segmentation, was generated based on merged fluorescent channels. The second neural network, the DNA fiber classification NN, was created using individual fluorescent channels to classify DNA fibers based on their red and green segments. To generate NNs, we utilized an interactive training module in the CellSens software to manually annotate hundreds of DNA fibers, followed by deep learning training to develop an initial AI model. This model was subsequently imported into the scanR analysis software to create ground truth. Afterward, this ground truth was used to train final NNs on a comprehensive dataset containing images with thousands of DNA fibers. After multiple rounds of validation, a scanR analysis assay was established to automatically analyze DNA fibers (Fig. 1d).

The first step in DNA fiber analysis involves segmenting DNA fibers using detection and classification NNs to segment whole DNA fibers as main objects or red and green DNA fiber parts as sub-objects, respectively (Fig. 1e). To optimize the segmentation of both main- and sub-objects, users can define intensity thresholds and set minimum and maximum object sizes. DNA fibers prepared using the spreading method (Fig. 1b) have a non-uniform distribution, leading to areas either devoid of DNA fibers or where fibers are densely packed. The latter poses challenges during the segmentation. To mitigate these issues, we have implemented two quality control (QC) parameters (Fig. 1f, g). The first parameter is based on the elongation factor, while the second, referred to as QC periphery, relies on modified object boundaries. These parameters, applied through the gating strategy, help to filter high-density DNA fiber regions and eliminate false-positive elongated fibers, ensuring the analysis of well-separated DNA fibers. Of note, when DNA fibers are prepared using the DNA combing technique, the application of QC parameters 1 and 2 can be disregarded, as the produced fibers inherently show sufficient separation. Optionally, unfocused, weak, or disconnected DNA fibers may be filtered out using QC parameter 3 based on the projection of the mean intensity of merged fluorescent channels (Fig. 3h), aiding the selection of high-quality DNA fibers in analysis. After applying QC parameters, DNA fibers are classified into five categories through the recognition of red and green segments using the classification NN. The classification criteria for red-green labeling (Fig. 1a) are as follows: ongoing fork (red and green segments), origin firing (green segment only), terminated/stalled fork (red segment only), bidirectional fork (red segment flanked by two green segments), and termination (green segment flanked by two red segments) (Fig. 1i). The multiparameter analysis framework enables a comprehensive examination of replication fork dynamics, incorporating parameters such as replication fork speed, symmetry, and the frequency of various DNA fiber classes (Fig. 1j), in conjunction with the aforementioned QC parameters. Finally, unbiased image galleries can be generated for individual DNA fiber classes (Fig. 1i), thereby enhancing the visual inspection of the analyzed DNA fibers. In summary, our qAID fiber workflow integrates automated imaging, segmentation, multiparameter analysis, and visual inspection of DNA fibers in a rapid and quantitative manner, yielding valuable insights into replication dynamics.

**Figure 3:**
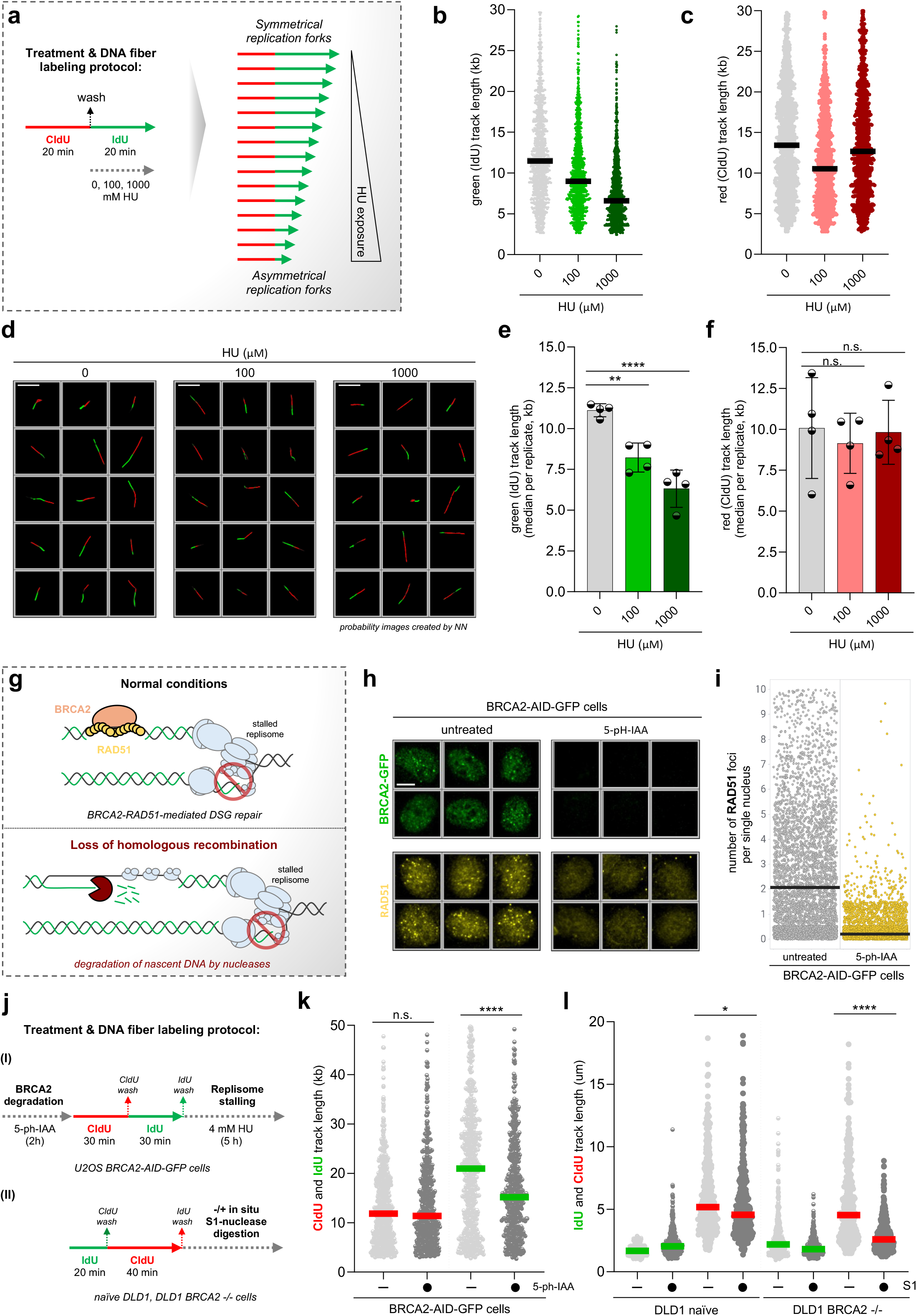
Quantitative analysis of replication fork response to low and high HU concentrations. **(a)** A schematic representation illustrating the DNA fiber labeling protocol to assess the replisome response to low and high HU concentrations. HU was added along with IdU, leading to the generation of asymmetrical replication forks. **(b)** qAID-based quantification of green track length of ongoing forks in U2OS cells treated with indicated HU concentrations. DNA fibers were prepared by spreading technique. Lines represent medians; n ≈ 1000 fibers per condition. **(c)** qAID-based quantification of red track length of ongoing forks in U2OS cells treated with indicated HU concentrations. DNA fibers were prepared by spreading technique. Lines represent medians; n ≈ 1000 fibers per condition. **(d)** Unbiased qAID galleries of DNA fibers in **(b, c)**. Probability images created by the classification network are displayed. Scale bar, 10 μm. **(e)** Quantification of qAID-based green track length plots in **(b)**; data are mean ± s.d.; n = 4 independent experiments. P values were calculated by ordinary one-way ANOVA with Dunnett’s test **P=0.0020, ****P < 0.0001. **(f)** Quantification of qAID-based red track length plots in **(c)**; data are mean ± s.d.; n = 4 independent experiments. P values were calculated by ordinary one-way ANOVA with Dunnett’s test. Not significant (n.s.) denotes P > 0.1. **(g)** A schematic representation illustrating the role of BRCA2 and RAD51 in the protection of nascent DNA to counteract its degradation by specific nucleases. **(h)** Unbiased QIBC galleries of BRCA2-AID-GFP U2OS cells immunostained for RAD51 upon indicated treatment (1 µM 5-ph-IAA for 2 h). Scale bar, 10 μm. **(i)** QIBC plots of BRCA2-AID-GFP U2OS cells immunostained for RAD51 upon indicated treatment. n ≈ 6,000 cells per condition. **(j)** DNA fiber labeling protocol with indicated treatment to study nascent DNA degradation in the presence and absence of BRCA2. **(k)** qAID-based quantification of red and green track length of ongoing forks in BRCA2-AID-GFP U2OS cells upon indicated treatment. DNA fibers were prepared by spreading technique. Lines represent medians; n ≈ 600 fibers per condition. P values were calculated by a two-tailed unpaired *t*-test ****P<0.0001, not significant (n.s.) denotes P > 0.1. **(l)** qAID-based quantification of red and green track length of ongoing forks in DLD1 naïve and BRCA2-knockout cells upon indicated S1 treatment. DNA fibers were prepared by spreading technique. Lines represent medians; n ≈ 400 fibers per condition. P values were calculated by a two-tailed unpaired *t*-test *P=0.0124, ****P<0.0001.

### The interplay between origin activation and the movement of individual replisomes

Accurate and complete DNA replication relies on the frequency of origin activation and the movement of individual replication forks along the DNA template. Recently, replication fork speed control was identified as a critical genome surveillance mechanism that safeguards replicating genomes from the escalation of physiological replication stress into more severe forms^13,14,22^. This mechanism facilitates the adaptation of the replication fork velocity in response to the dynamic cellular environment, including fluctuations in the origin firing level^23,24^. Specifically, the frequency of origin activation inversely affects the replication fork speed (Fig. 2a). To quantitatively investigate the interplay between origin activation and the movement of individual replisomes, we modulate the levels of origin activation by ATR and CDC7 inhibitors, which increase and decrease the conversion of inactive MCMs to active replisomes, respectively (Fig. 2a). First, we employed QIBC to assess DNA replication globally at the single-cell resolution^25^. While chromatin binding of replisome components, such as CDC45 or TIMELESS, increased approximately four-fold in the presence of ATR inhibitor (Fig. 2b, Extended Data Fig. 3a-g), PCNA binding showed only a modest increase following ATR inhibitor treatment (Extended Data Fig. 4a-d). Importantly, this modest increase in PCNA binding on chromatin correlated with the level of DNA synthesis measured by QIBC of EdU incorporation at the single-cell level (Fig. 2c; Extended Data Fig. 4e-g) as well as with the level of origin firing at the single replication fork resolution measured by qAID (Fig. 2d, e). These observations are consistent with recent pre-prints^26-28^, demonstrating that the natural depletion of PAF15 under replication stress conditions leads to the exhaustion of functional PCNA at the lagging strand. This deficiency ultimately causes RPA exhaustion and replication catastrophe^11,29^. Notably, all ATR inhibition-induced phenotypes were reversed by concurrent CDC7 inhibitor treatment, which reduces origin firing.

At the single replisome level, ATR inhibition led to a decreased frequency of ongoing replication forks (Extended Data Fig. 5a) and a dramatic reduction in replication fork speed analyzed through various approaches (Fig. 2f-l; Extended Data Fig. 5b). In detail, manual measurements of the lengths of 350 fibers demonstrated that the movement of individual replication forks was dramatically diminished following ATR inhibitor treatment; however, this replication fork slowdown was restored with a CDC7 inhibitor (Fig. 2f). The same trend was observed during the automated analysis of DNA fibers using qAID workflow, which enabled the analysis of approximately one thousand DNA fibers (Fig. 2g) and allowed their visual inspection through unbiased image galleries (Fig. 2h). The observed phenotypes were consistent across biological replicates of the DNA fiber datasets analyzed via both manual and automated approaches (Fig. 2i, j). Importantly, the automated analysis of the DNA fiber dataset prepared by DNA combing in the Moudry laboratory aligned with observations from DNA fiber spreading (Fig. 2k, l). These observations reinforce previous conclusions that individual replisomes can adjust their movement in response to origin firing levels ^23,24^. Furthermore, our comprehensive analysis has verified that the qAID workflow is a reliable and robust method for automated analysis of DNA fibers, irrespective of the variations in preparation methods across different laboratories.

To demonstrate the effectiveness of the qAID workflow, we performed a comprehensive DNA fiber analysis on a panel of cell lines, including both non-cancerous and cancerous samples (Extended Data Fig. 5c-e). Our findings confirmed a previously observed phenomenon—non-cancerous cells displayed faster replication fork progression, whereas cancer cells exhibited slower replication fork movement (Extended Data Fig. 5c-e)^14^. Decreased replication fork progression in cancer cells may be attributed to the complex intrinsic properties of cancer cells, including variations in origin firing and metabolic activity^14^. In summary, we conclude that qAID workflow serves as a reliable tool to automatically measure the frequency of DNA fiber classes and replication fork speed. Integrating qAID with QIBC provides a powerful methodology for investigating DNA replication dynamics and its alteration at both global and local scales.

### DNA replication fork response to mild and high replication stress induced by hydroxyurea

To explore how replisome machinery responds to varying levels of RS across different genetic backgrounds, we treated cells with a range of hydroxyurea (HU) concentrations (Fig. 3a). HU inhibits ribonucleotide reductase (RNR), the enzyme responsible for the conversion of ribonucleotides into deoxyribonucleotides (dNTPs)^30^. At high concentrations, HU substantially depletes dNTPs, resulting in replication fork stalling. Conversely, at lower concentrations of HU, no significant alterations in dNTP levels were observed^14^. Instead, the replication fork slowdown under these conditions is driven by redox imbalance and the release of reactive oxygen species (ROS). These changes are detected by PRDX2, a replisome-based redox sensor, which modulates the replication fork speed in accordance with the nuclear redox state^14^. To quantitatively assess this replisome response, we treated cells with low and high HU concentrations during the second labeling step of DNA fiber protocol, which will lead to the generation of asymmetrical replication forks (Fig. 3a). Aligned with these expectations, qAID analysis revealed a gradual decrease in replication fork speed that correlated with the HU concentration (Extended Data Fig. 6a, b). Examination of individual DNA fiber segments revealed a shortening of the green tracks, while the red tracks remained constant (Fig. 3b, c). These observations were further validated through visual inspection of unbiased DNA fiber galleries (Fig. 3d) and were consistent across the biological replicates (Fig. 3e, f). Additionally, these phenotypes were confirmed by the QIBC-based monitoring of the DNA replication program at the single-cell level (Extended Data Fig. 6c-i; Extended Data Fig. 7a).

Acute HU treatment has been widely used to investigate the molecular mechanism that protects nascent DNA^9^. Beyond their well-established roles in homologous recombination (HR)-mediated repair of double-strand breaks, BRCA2 and RAD51 are also crucial for preserving nascent DNA by preventing its degradation by nucleases such as MRE11 and EXO1 (Fig. 3g)^31-35^. To examine the effect of HU on nascent DNA stability in HR-deficient conditions, we generated a BRCA2 auxin-inducible degron in U2OS cells (Extended Data Fig. 7b, c), enabling rapid degradation of BRCA2 without cell cycle alterations (Extended Data Fig. 7d-g). Since BRCA2 is instrumental in facilitating RAD51 recruitment to single-stranded DNA therefore, its rapid depletion considerably impaired RAD51 loading on nascent DNA (Fig. 3h, i; Extended Data Fig. 7h). Under these HR-deficient conditions, we performed DNA fiber assay in the presence of HU to analyze the stability of nascent DNA strands by qAID (Fig. 3j (I)). The automated analysis revealed a shortening of the green DNA fiber segments, indicating the degradation of nascent DNA in the BRCA2-deficient cells (Fig. 3k; Extended Data Fig. 7 i, j). Recently, several studies have demonstrated the presence of daughter strand gaps (DSGs) in newly synthesized DNA within BRCA2-depleted cells^7-9^. These DSGs can be detected using electron microscopy^36,37^, super-resolution microscopy^38^, or a modified DNA fiber assay with S1 nuclease digestion (Fig. 3j (II))^39-41^. To quantitatively assess DSGs in normal and HR-deficient cells, we employed a well-established DLD1 cellular model, which includes both wild-type and BRCA2-knock out cells and performed a DNA fiber assay with S1 nuclease treatment (Fig. 3j (II); Extended Data Fig. 7k). Consistent with previous research^41,42^, we observed a substantial shortening of the IdU-labeled DNA tracts after S1 nuclease treatment, confirming the presence of DSGs (Fig. 3l). Overall, our findings quantitatively validate previous observations on DNA replication fork responses to varying levels of HU-induced RS. Furthermore, we demonstrate that qAID is a powerful tool for detecting subtle replication fork asymmetries, particularly those associated with nascent DNA degradation in HR-deficient cells.

## DISCUSSION

Replication stress is a major driver of genome instability. Understanding its molecular mechanisms through quantitative and unbiased methodologies could facilitate early diagnosis and the development of effective therapies for diseases associated with genomic instability^3,4^. QIBC, which employs the scanR high-content imaging system, has been widely used to study RS at the single-cell level^11-15^. Decades of research have established the DNA fiber method as a powerful tool for exploring replication fork dynamics and the mechanisms underlying RS at the single replication fork resolution^16-18^. However, unlike single-cell RS analysis, DNA fiber assays depend heavily on manual acquisition and measurement of replication fork parameters (Extended Data Fig. 1c). This manual approach is time-consuming and prone to user bias, potentially affecting result interpretation and reproducibility. To address these challenges, we developed qAID—an automated workflow that integrates the scanR high-content imaging system with the TruAI deep learning module for DNA fiber acquisition and analysis.

The qAID workflow allows imaging and multiparameter analysis of thousands of DNA fibers within a timeframe of several dozen minutes (Fig. 1). The multiparameter analysis of DNA fibers contains three main features: 1) Automated DNA fiber segmentation enabling the detection of whole DNA fibers along with their corresponding red and green segments utilizing two neural networks (Fig. 1e). 2) Quality control parameters integrated into a gating strategy to ensure the selection of high-quality fibers based on predefined criteria (Fig. 1f-h). 3) Examination of replication fork dynamics, including replication fork speed, symmetry, and the frequency of various DNA fiber classes (Fig. 1j). Users can customize thresholds for each feature, including intensity levels and object size limits. This setup retains the flexibility to integrate variations in DNA fiber preparation across different laboratories while enhancing the result reproducibility. Although DNA fibers can be automatically captured using a scanR high-content imaging system, this is not a requirement for conducting multiparameter analysis. DNA fiber images from various imaging systems can be imported into the scanR analysis software. We validated this capability by analyzing DNA fibers from multiple imaging systems beyond scanR. The NNs developed in this study were trained on DNA fiber datasets from the scanR high-content imaging system (Evident) and the Axio Imager microscope (Zeiss) with a pixel size of 0.16125 μm/pixel. The multiparameter analysis settings enable flexible scaling of the NNs, allowing adaptation to DNA fibers captured at different resolutions (Extended Data Fig. 1b). Moreover, users may use the NNs developed in this study to train own NNs on their samples to improve DNA fiber segmentation or generate new models without requiring extensive programming expertise. Additionally, multiparameter analysis includes automatic generation of unbiased image galleries for enhanced visual inspection of analyzed DNA fibers (Fig. 1i). We believe that the qAID workflow enhances DNA fiber analysis efficiency and improves the result reproducibility.

Several attempts have been made to develop software for the automated analysis of DNA fibers^43-46^. Although these tools are freely available, none of them have been widely adopted, as shown by the analysis of 110 studies using DNA fibers (Extended Data Fig. 1c). The limited adoption is largely attributed to the lack of flexibility in DNA fiber analysis, including restricted parameter adjustments, the inability to train customized networks, and insufficient validation across diverse experimental settings. In this study, we developed a tool that enables users to customize the qAID workflow to their specific requirements while providing robust validation of its functionality. Our results demonstrate that qAID reliably measures the frequency of DNA fiber classes, replication fork speed, and subtle variations in fork symmetry, particularly in relation to nascent DNA degradation in HR-deficient cells (Fig. 2, 3). Our methodology was validated by analyzing DNA fibers prepared using both spreading and combing techniques in three independent laboratories. Although the workflow is based on scanR software, all assay files and NNs are freely available and may serve as a foundation for future advancements, such as extending qAID to automated analysis of origin distances or dynamics of replication forks at defined genomic loci. In conclusion, the qAID workflow provides a rapid and efficient method for examining replication dynamics and its alterations at a single-molecule resolution. This innovative AI-based method paves the way for a deeper understanding of the molecular mechanisms underlying RS and its association with severe diseases of human health.

## ACKNOWLEDGMENTS

The research work in the Sedlackova laboratory was supported by the Czech Science Foundation Junior Star (grant no. 22-20303M), the European Union’s Horizon 2022 Widera Talent program (ERA grant agreement no. 101090292), EMBO Installation Grant (grant no. IG-5689-2024), and the Czechoslovak Microscopy Society stipendiary. The research leading to these results in Costanzo’s lab received funding from AIRC under IG 2023—ID. 28725 project. A.A. was supported by the 31406 AIRC fellowship for Italy. The work in the Moudry lab was supported by the MEYS CR (Large RI Project LM2023050LM2018129 -Czech-BioImaging) and the SALVAGE project, registration number: CZ.02.01.01/00/22_008/0004644, supported by OP JAK, with co-financing from the EU and the State Budget.

We sincerely thank Jiri Polasek, Tomas Pop, and Tomas Jendrulek for the technical maintenance of high-content imaging microscopes. We are also deeply grateful to Maj-Britt Rask and Claudia Lukas for their support and for generously providing essential reagents. We would like to extend our special thanks to Manoel Veiga from Evident for the critical reading of the manuscript.

## AUTHOR CONTRIBUTION

P.F. validated the qAID workflow, performed QIBC and DNA fiber spreading experiments, analyzed the data, and contributed to experimental design and manuscript writing. T.D. trained neural networks for DNA fiber detection and, together with H.P.-S. developed qAID workflow. A.A. performed DNA fibers treated with S1 nuclease. Z.M. performed DNA combing experiments. S.N. contributed to HU experiments. A.K.Y. generated BRCA2 degron cell line. P.M. supervised Z.M. and contributed to manuscript editing. V.C. supervised A.A. and contributed to manuscript editing. H.P.-S. conceived the project, contributed to cell line generation, designed experiments, analyzed data, prepared figures, and wrote the manuscript. All authors read and commented on the manuscript.

## MATERIAL & CORRESPONDENCE

Should be addressed to H.P.-S.

## METHODS

### Cell Culture

The human U2OS osteosarcoma cell line, MCF7 adenocarcinoma epithelial cell line, HCT 116 colorectal carcinoma cell line, breast cancer CAL51 cell line, primary foreskin BJ fibroblasts, and primary retinal epithelial RPE-1-hTERT cell line were grown under standard conditions (37°C and 5% CO_2_) in Dulbecco’s modified Eagle’s medium (DMEM) containing high glucose and GlutaMAX (Thermo Fisher Scientific, 31966047), and supplemented with 10% fetal bovine serum (FBS; Thermo Fisher Scientific, A5256801) together with 0.5% penicillin-streptomycin (Thermo Fisher Scientific, 15140122). A derivative of the U2OS cell line expressing C-terminally endogenously GFP-tagged CDC45 was generated using CRISPR-Cas9 and validated in the previous study^15^. DLD-1 wild-type (Cat#HD PAR-008; RRID:CVCL_0248) and DLD-1 BRCA2-deficient (Cat#HD 105-007; RRID:CVCL_HD57) cells were acquired from Horizon discovery. Cells were cultured in RPMI 1640 (Lonza) supplemented with 2 mM glutamine, 10% Fetal bovine serum (FBS), and 1% penicillin-streptomycin (Pen-Strep). Cells were maintained at 37°C and 5% CO_2_. All cell lines and their derivatives were routinely tested for the absence of mycoplasma contamination using the Mycoplasma Detection Kit (InvivoGen, rep-mys-50).

### Generation of BRCA2 degron cell line

U2OS cell line expressing C-terminally endogenously AID-GFP tagged BRCA2 protein was generated using CRISPR-Cas9 as previously described^15^. Briefly, single-guide RNA (gRNA) targeting the C-terminal of the BRCA2 locus (ATATATCTAAGCATTTGCAA) was cloned into the pX330-U6-Chimeric_BB-CBh-hSpCas9 vector (Addgene plasmid 42230, a gift from F. Zhang) via the *Bbs*I restriction site. U2OS cells were co-transfected with a pX330 plasmid containing cloned gRNA and a donor plasmid containing the tag (AID-GFP) with a flexible linker flanked by 900 bp homology arms complementary to the C-terminus of BRCA2. Transfection was carried out using Lipofectamine LTX with Plus reagent (Thermo Fisher Scientific, 15338-100). Transfected cells were grown for 8 days followed by cell sorting to isolate GFP-positive cells. After 5 days, the sorted cells were serially diluted into 100 mm dishes to obtain single colonies, which were expanded for further characterization by western blotting and junction PCR spanning the C-terminal of BRCA2. Selected clones were functionally validated by immunofluorescence using QIBC. Only the cell lines that passed all validation steps were used for further experiments. Clones with homozygous endogenous tagging were transfected with degron 2 plasmid (pCMV6-A-puro-TIR1(F74G)-9xMyc) containing a codon-optimized (specific for human) TIR1 gene^47^. Transfected cells were serially diluted into 100 mm dishes and selected with DMEM media containing geneticin (G418, 0.4 mg/ml, Merck, 472787001) for 10 days to obtain single colonies. After clone picking, colonies were expanded and tested for TIR1 expression by immunofluorescence using an antibody against the Myc tag. BRCA2 degradation was achieved by 5-ph-IAA (Merck, SML3574) at a final concentration of 1 μM.

### Chemical reagents and antibodies

ATR inhibitor (AZD6738, Selleckchem, S7693), CDC7 inhibitor (BMS-863233, Selleckchem, S7547), hydroxyurea (Merck, H8627), 5-pH-IAA degron (Merck, SML3574), 5-ethynyl-2’-deoxyuridine (EdU, Thermo Fisher Scientific, C10640), Alexa Fluor 647 picolyl azide (Thermo Fisher Scientific, C10640) for EdU Click-IT reaction, 5-Chloro-2′-deoxyuridine (CldU, Merck, C6891), 5-Iodo-2′-deoxyuridine (IdU, Merck, 17125), and S1 nuclease (Thermo Fisher Scientific, ENO321) were used as indicated in corresponding figure legends. For immunofluorescence, the following primary antibodies were used: GFP (rabbit, Chromotek, PABG1, 1:1000), TIMELESS (rabbit, Abcam, ab109512, 1:1000), PCNA (human, Immuno Concepts, 2037, 1:1000), RAD51 (rabbit, BioAcademia, 70-012, 1:1000). Secondary antibodies for immunofluorescence: goat anti-rabbit Alexa Fluor 568 (Thermo Fisher Scientific, A-11036, 1:2000), goat anti-rabbit Alexa Fluor 488 (Thermo Fisher Scientific, A-11034, 1:2000), and donkey anti-human Alexa Fluor 647 (Jackson Immuno Research, 709-605-149, 1:2000). For Western Blot, the primary antibodies used were BRCA2 (mouse, Merck, OP95, 1:1000), MCM7 (mouse, Santa Cruz, sc-9966, 1:1000), GFP (rabbit, Chromotek, PABG1, 1:1000), and vinculin (mouse, Merck, V9131, 1:20000). For Western Blot, HRP-conjugated anti-mouse (Vector Laboratories, PI-2000, 1:10000) and anti-rabbit (Vector Laboratories, PI-1000-1, 1:10000) secondary antibodies were used.

### DNA fiber spreading

Approximately 100,000 cells were seeded per well (35 mm) a day before the DNA fiber experiment. The next day, cells were subjected to specific treatment as detailed in protocols attached in figure panels and labeled with thymidine analogs (25 µM CldU and 250 µM IdU), separated by three washes with phosphate-buffered saline 1x (PBS). Subsequently, labeled cells were collected in ice-cold PBS and mixed with unlabelled cells in a 1:1 ratio. To lysate the cells, 4 µl of cell suspension was placed on a positively charged slide (Menzel Gläser, SuperFrost, VWR, 631-0705) and mixed with 8 µl of lysis buffer (0.5 % SDS, 200 mM Tris pH 7.5, 50 mM EDTA) by thorough pipetting. After 2 minutes of *in situ* cell lysis, slides were inclined at an angle of approximately 15° to facilitate DNA fiber spreading. Then, DNA spreads were left to be air-dried briefly. Next, slides were fixed in methanol:acetic acid (3:1) solution for 10 minutes at the laboratory temperature and washed three times in PBS. After fixation, DNA fibers were denatured at 2.5 M HCl for 80 minutes at the laboratory temperature. Next, denatured DNA fibers were neutralized by washing the slides four times in PBS and then blocked with a blocking buffer solution (1x PBS 0.1% TritonX-100, 1% BSA) for 30 minutes. After blocking, to detect CldU, slides were incubated with rat monoclonal anti-BrdU antibody (ab6326, Abcam, 1:100 in blocking buffer) for 75 minutes, washed once with 0.1% Tween PBS and twice with PBS, fixed in 4% paraformaldehyde for 10 minutes, and further incubated with the secondary donkey anti-rat Alexa Fluor 594 (Thermo Fisher Scientific, A21209, 1:200) for 60 minutes. Slides were washed thrice with PBS and incubated with mouse anti-BrdU (Becton Dickinson, 347580, 1:100) overnight at 4 °C to allow the detection of IdU. The next day, slides were washed once with 0.1% Tween PBS 1x and twice with PBS and incubated with the secondary anti-mouse Alexa Fluor 488 (Thermo Fisher Scientific, A-11029, 1:200) for 60 minutes. After washing three times with PBS, slides were air-dried and mounted with a Mowiol-based mounting medium (12% Mowiol 4-88 (Merck, 81381-250G), 30% glycerol, 0.12 M Tris-HCl pH 8.5).

### DNA fiber combing

DNA combing was performed as previously described^48^. Approximately 400,000 cells were seeded per well (60 mm) two days before the DNA fiber experiment. Exponentially growing cells were labeled with 25 µM CldU for 20 minutes, washed with DMEM, and then labeled with 250 µM IdU for 20 minutes, either with 5 µM ATR inhibitor alone, in combination with 5 µM CDC7 inhibitor or without inhibitors. Replicating cells were halted and collected in ice-cold PBS, and DNA was extracted using a FiberPrep kit (Genomic Vision, EXT-001) following the manufacturer’s instructions. Once extracted, DNA was combed on vinylsilane-coated CombiCoverslips (Genomic Vision, COV-002-RUO) with a constant speed of 0.3 mm/s. DNA was then baked for 2 hours at 65°C and stored at -20°C. For immunostaining, combed DNA was denatured with buffer D (0.5 M NaOH, 1M NaCl) for 8 minutes, dehydrated with 70%, 90%, and 100% ethanol washes, and blocked using ADM buffer (10% FBS in DMEM) for 30 minutes. Coverslips containing DNA fibers were immunostained with mouse anti-BrdU (BD Biosciences, BD347580, 1:10) and rat anti-BrdU (Abcam, ab6326, 1:50) primary antibodies for 1 hour in ADM buffer. After four washes with PBS, coverslips were incubated with anti-mouse Alexa Fluor 488 (Invitrogen, A11001, 1:100) and goat anti-rat Alexa Fluor 568 (Abcam, ab175476, 1:100) secondary antibodies for 1 hour in ADM buffer. After four washes with PBS, coverslips were air-dried and mounted using Vectashield (Vector Laboratories, H1000).

### DNA fiber assay with S1 nuclease

Both DLD1 wild-type and BRCA2-deficient cells were grown in 6-well plates (450,000 cells per well) for 24 hours and pulse-labelled with 50 µM CldU (Merck) for 20 minutes and with 250 µM IdU (Merck) for an additional 40 minutes. Thereafter, cells were trypsinized, counted, and resuspended in CSK100 buffer (100 mM NaCl, 10 mM PIPES pH 6.8, 3 mM MgCl_2_, 300 mM sucrose, 0.5% Triton X-100). Next, cells were permeabilized for 5 minutes at room temperature, centrifuged for 5 minutes at 7000 rpm, and resuspended in 1 mL of S1 buffer (30 mM Sodium acetate pH 4, 2 mM Zinc sulfate, 50 mM NaCl, 5% glycerol). The nuclei were incubated for 30 minutes at 37°C with 10 U/mL of S1 nuclease (Sigma) in S1 buffer and then centrifuged for 5 minutes at 7000 rpm and resuspended in PBS to a final concentration of 1000 nuclei/µL. Two microliters of the cell suspension were lysed on a clean glass slide with 8/9 µL of MES lysis buffer (50 mM MES pH 5.6, 0.5% SDS, 50 mM EDTA) for 6 minutes. The slide was tilted at a 15-degree angle, air-dried for 30 minutes, and fixed in freshly prepared acetic acid/methanol (1:3) for 10 minutes, and then air-dried and stored at 4°C overnight. The following day, the slides were rehydrated with 1x PBS for 5 minutes, and the DNA was denatured with 2.5 M HCl for 60 minutes, followed by two washes of 5 minutes with PBS. Next, the slides were then blocked in 3% BSA for 30 minutes, followed by incubation with a primary antibodies Rat anti-BrdU (Abcam, 6326) and mouse anti-BrdU (BD Biosciences), both at a dilution of 1:50, in blocking solution for 90 minutes at 37°C in a humid chamber. After incubation, the slides were washed 3 times with PBS for 5 minutes each and incubated with secondary antibodies donkey anti-mouse Alexa 568 (1:150, Thermo Fisher) and chicken anti-rat Alexa 488 (1:150, Thermo Fisher) in blocking solution for 45 minutes at 37°C in a humid chamber. The slides were then washed three times with PBS for 5 minutes each, air-dried, mounted in Vectashield plus (Vector Labs), and stored at -20°C until image acquisition.

### Automated DNA fiber acquisition

DNA fiber images for DNA spreading and S1 nuclease experiments were acquired using a scanR high-content imaging system equipped with IX83 microscope frame (Evident), UPLXAPO dry objective (40x, 0.95 NA), fast excitation and emission filter wheel for DAPI, FITC, Cy3, and Cy5 wavelengths, Spectra X led fluorescence light source (Lumencor), digital monochromatic sCMOS ORCA-Flash 4.0 LT Plus camera (Hamamatsu, resolution: 2048×2048, pixel size: 6.5×6.5 μm, effective area: 13.312×13.312 mm) and Z-Drift Compensation System IX-ZDC. The microscopy system was controlled through the scanR acquisition software (Evident, v3.5.1), in which autofocus settings, acquisition settings for individual fluorescent channels, and scanning patterns were defined. To set up the autofocus, the Z-offset was calibrated, and the refractive index was adjusted to calculate the thickness value. The parameters of autofocus were as follows: green LED excitation with 100% intensity, Cy3 emission filter, 40x UPLXAPO dry objective, 100 ms exposure time, and 2×2 camera binning. For two color DNA fibers labeled with Alexa Fluor 488 and 594 (or 568) dyes, FITC and Cy3 fluorescent channels with appropriate filter cubes were used. FITC channel was defined as follows: cyan LED excitation with 100% intensity, FITC emission filter, 40x UPLXAPO dry objective, 1000 ms exposure time, no pre-acquisition delay, frame averaging 1, Z-stack layers 1, and 1×1 camera binning. Cy3 channel was defined as follows: green LED excitation with 100% intensity, Cy3 emission filter, 40x UPLXAPO dry objective, 1000 ms exposure time, no pre-acquisition delay, frame averaging 1, Z-stack layers 1, and 1×1 camera binning. Due to the uneven distribution of DNA fibers across the slide when employing the DNA fiber spreading technique (Fig. 1b), we typically select 6-8 locations across the slide using the pattern position wizard. For each location, we conducted automated scans of 9 to 16 positions, yielding the collection of approximately one thousand DNA fibers per sample.

DNA fiber images for DNA combing were acquired using a scanR high-content imaging system equipped with IX81 microscope frame (Evident), UPlanSApo dry objective (40x, 0.95 NA), fast excitation, and emission filter wheel for DAPI, FITC, Cy3, and Cy5 wavelengths, MT20 illumination system and digital monochromatic sCMOS ORCA-Fusion C14440-20UP camera (Hamamatsu, resolution: 2304×2304, pixel size: 6.5×6.5 μm, effective area: 14.976×14.976 mm). The microscopy system was controlled through the scanR acquisition software (Evident, v3.3.0), in which autofocus settings and acquisition settings for individual fluorescent channels were defined as described above. Since DNA fibers are uniformly distributed when employing the DNA fiber combing technique (Fig. 1b), we typically select one center location on the coverslip and conduct automated scans of 49-64 positions, yielding the collection of approximately one thousand DNA fibers per sample. Further details on DNA fiber acquisition using the scanR high-content imaging system are provided in the step-by-step user guide (Supplementary File 1).

### Automated DNA fiber analysis

To develop NNs detecting and classifying DNA fibers, we employed the TruAI deep learning module in CellSens software (Evident, v4.2.1) and scanR analysis software (Evident, v3.4.1.). The DNA fiber detection NN dedicated to whole-fiber segmentation was generated based on merged fluorescent channels. The DNA fiber classification NN was created using individual fluorescent channels to classify DNA fibers based on their red and green fiber segments. First, DNA fibers were manually annotated using an interactive training module in the CellSens. The initial fiber dataset contained 19 images, each with approximately 20-50 fibers acquired in the scanR high-content imaging systems equipped with IX83 and IX81 microscope frames (Evident) and the Axio Imager microscope (ZEISS) with a pixel size of 0.16125 μm/pixel. The manual annotation was followed by deep learning training in CellSens software using image segmentation and Specific Network (network architecture: U-Net; data sampler: balanced; normalization: image standardization; loss: softmax cross entropy; optimizer: adam; geometric augmentation: 90°rotations, mirroring; image processing augmentation: none) as a training configuration. This led to the development of the initial AI model based on the larger DNA fiber dataset, containing 68 images in total. Next, this AI model was imported into the scanR software to create a ground truth, which was curated by gating to filter out false positive fiber detection. The training of final networks was done by neural network training in scanR software using semantic segmentation and Specific Network (network architecture: U-Net; data sampler: balanced; normalization: image standardization; loss: softmax cross entropy; optimizer: adam; geometric augmentation: 90°rotations, mirroring; image processing augmentation: none) as a training configuration, applied on a comprehensive dataset (1772 DNA fiber images in total). The final DNA fiber detection NN was created after reaching 25,000 iterations and 0.901 similarity (value evaluating prediction probability to manual annotation), and the final DNA fiber classification NN was created after reaching 500,000 iterations and 0.784 similarity. The validation of NNs was done by manual error scoring of DNA fiber detection or classification on the 100 random examples per each fiber category. Depending on the nature of the spotted errors, we adjusted the manual DNA fiber annotation in CellSens or curated the gating of ground truth in scanR and retrained the network to achieve a success score for all DNA fiber categories of around 90% or higher.

After multiple rounds of validation, we established a scanR-based assay with three main features: automated DNA fiber segmentation, quality control parameters, and measurement of replication fork dynamics. To segment the whole DNA fiber as the main object, DNA fiber detection NN was used with the following settings: 30,000 intensity threshold, 50 pixels minimum object size, and 10,000 pixels maximum object size. To segment red and green DNA fiber sub-objects, DNA classification NN was used with the following settings for each color: 15,000 intensity threshold, 15 pixels minimum object size, and 10,000 pixels maximum object size. In the assay settings, we specified the core parameters for DNA fiber analysis, including the elongation factor for QC parameter 1, QC periphery for QC parameter 2 (based on modified object boundaries with settings distance: 2, width: 5, overlap treatment: segment), mean intensity of red, green and merged channels for QC parameter 3, red and green segments counts for classification of DNA fibers, perimeter and max Feret Diameter to determine DNA fiber length. All described parameters were defined for individual fluorescent channels within the main object or corresponding sub-objects. DNA fiber length in pixels can be calculated through the standard perimeter formula or max Ferret Diameter; both methods yield the same result (further details and direct comparisons are provided in Supplementary File 1). The formulas for direct calculations of DNA fiber length in μm and kb or replication fork speed were integrated into the scanR assay as derived parameters with conversion factors 0.16125 μm per pixel and 2.59 μm per kb with respect to labeling time. Further refinement of QC parameters 1-3 was achieved through a specific gating strategy as outlined in Fig. 1f-h. Tightening or loosening specific gates was applied to achieve a minimum of 90% DNA fiber classification success for each DNA fiber category. The multiparameter analysis was exported as a table and further analyzed in Spotfire software (TIBCO, v.11.8.0) or GraphPad Prism (10.1.2.). Further details on DNA fiber analysis using the scanR software and step-by-step user guide are provided in Supplementary File 1.

### Immunofluorescence staining

Cells were plated in 6-wells containing 12 mm diameter, 1.5 mm-thick glass coverslips (VWR, MENZCB00120RAC20) and let to grow till 70-80% confluency the day of the experiment. To study chromatin-bound proteins, cells were pre-extracted with ice-cold cytoskeleton buffer (10 mM Hepes pH 7.5; 300 mM sucrose; 100 mM NaCl; 3 mM MgCl_2_ and 0.5% TritonX-100) for 10 minutes at room temperature, washed three times with PBS and fixed with 4% buffered formaldehyde (VWR, 9713.1000) for 20 minutes at room temperature. To stain the nuclear proteins, cells were fixed, omitting the pre-extraction step, with 4% buffered formaldehyde for 20 minutes at room temperature, washed thrice with PBS, and permeabilized with ice-cold PBS containing 0.2% TritonX-100 for 5 minutes at laboratory temperature. Primary antibodies were diluted in DMEM containing 10% FBS and incubated for 90 minutes at room temperature. For EdU detection, cells were incubated with Click iT buffer (100 mM Tris-HCl pH 8, 2 mM CuSO_4_, 1 ng Alexa Fluor 647 Azide, and 100 mM ascorbic acid) at room temperature for 30 minutes prior to primary antibody staining. Secondary antibodies together with 4’,6-Diamidino-2-Phenylindole, Dihydrochloride (DAPI, Thermo Fisher Scientific, D1306) were diluted in DMEM supplemented with 10% FBS and incubated with the coverslips for 45 minutes at laboratory temperature. Next, coverslips were washed thrice with PBS and twice with distilled water and mounted on slides using a Mowiol-based mounting medium.

### Quantitative image-based cytometry (QIBC)

Images for QIBC were acquired using a scanR high-content imaging system equipped with IX83 microscope frame (Evident), UPLXAPO dry objective (20x, 0.8 NA), fast excitation and emission filter wheel for DAPI, FITC, Cy3, and Cy5 wavelengths, Spectra X led fluorescence light source (Lumencor), digital monochromatic sCMOS ORCA-Flash 4.0 LT Plus camera, Z-Drift Compensation System IX-ZDC, and scanR acquisition software (Evident, v3.5.1). During image acquisition, the LED excitation intensity was set at 100%, and exposure times were adjusted individually for each fluorophore. Image analysis was performed in scanR analysis software (Evident, v.3.5.1), where an automated dynamic background correction was applied for each fluorescent channel separately and maintained the same for all images within a single experiment. Nuclei segmentation was performed through an intensity-threshold-based mask generated based on the DAPI signal. This mask was then applied to analyze pixel intensities in different channels for each nucleus. Count and fluorescent intensities for BRCA2 and RAD51 foci were measured using the Spot Detector tool as sub-objects. The multiparameter analysis was exported as a table and further analyzed in Spotfire software (TIBCO, v.11.8.0). A similar number of cells was analyzed and compared for each condition within the experiment. For visualization, low x-axis jittering was applied (random displacement of objects along the x-axis) to make overlapping objects visible.

### SDS-page and western blotting

Cell lysates were obtained by incubating cells with lysis buffer (10 mM Hepes pH 7.5, 500 nM NaCl, 1 mM EDTA, 1% NP-40) supplemented with protease and phosphatase inhibitors (ROCHE, 4693116001 and 4906845001) and 250 U per ml of benzonase (Merck, E1014-25KU). Cell lysates were denatured at 95°C for 5 minutes in Laemmli loading buffer containing LDS Sample Buffer (Thermo Fisher Scientific, NP0007) and NuPAGE Sample Reducing Agent (Thermo Fisher Scientific, NP0009), and separated on NuPAGE™ 3 to 8% Tris – Acetate precast gels (Thermo Fisher Scientific, EA03752BOX) following standard procedures. Separated proteins were then transferred to nitrocellulose membranes (Thermo Fisher Scientific, IB23002×3) using the iBlot system (Thermo Fisher Scientific, IB21001). After transfer, nitrocellulose membranes were incubated in a blocking buffer containing 1x PBS, 5% powdered milk (Merck, 70166-500G), and 0.1 % Tween-20 (Merck, P1379-500ML) for 1 hour at room temperature. Primary antibodies were diluted in a blocking buffer and incubated at 4 °C overnight, while secondary antibodies were diluted in a blocking buffer and incubated for 1 hour at room temperature. Protein signals were detected using ECL Select Western Blotting Detection Reagent (Merck, GERPN2235).

### Statistical analysis

Statistical analysis was performed using GraphPad Prism 10.1.2. Experiments were not randomized, and no blinding was used during data analysis. The sample size, statistical tests, and number of replicates for all experiments are specified in the figure legends.

### Data availability statement

All the source data, including numerical and statistical source data, will be deposited appropriately at the European Bioinformatics Institute (EBI) BioStudies database (https://www.ebi.ac.uk/biostudies/) and will be immediately accessible as an open resource pending acceptance of this manuscript. Any additional data or information in support of this study will be available from the corresponding author upon reasonable request.

