## Extended Data Figures 1-7 for "Quantitative AI-based DNA fiber workflow to study replication stress"

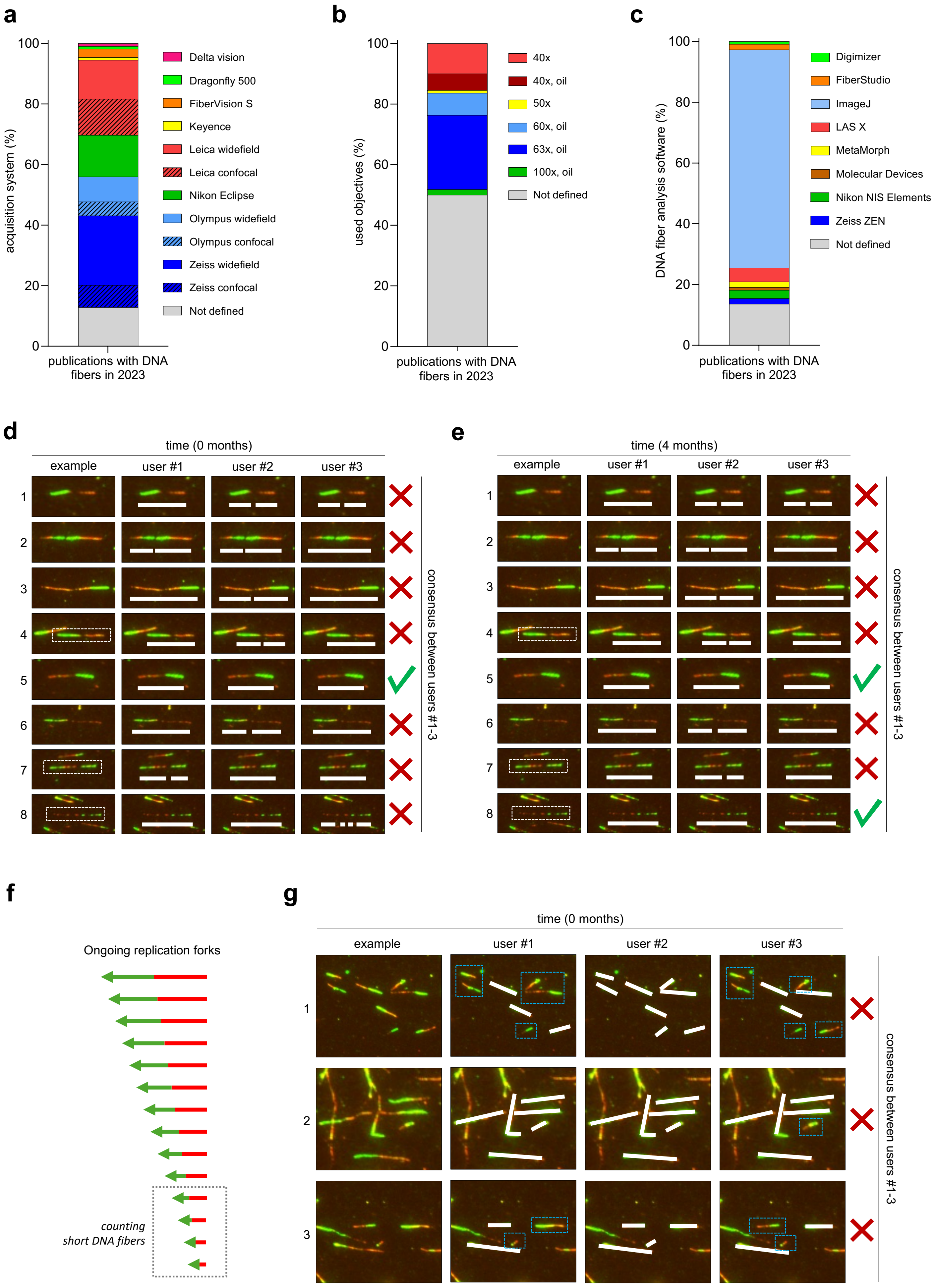


**Extended Data Figure 1**

**Extended Data Figure 1: Assessment of acquisition and analysis of DNA fibers in research papers published in 2023 and inter-user variability in manual DNA fiber segmentation. (a)** A bar chart illustrating variability in the DNA fiber acquisition systems in 110 PubMed publications with DNA fibers published in 2023. **(b)** A bar chart illustrating variability in the objectives to acquire DNA fibers in 110 PubMed publications with DNA fibers published in 2023. **(c)** A bar chart illustrating a predominant manual analysis of DNA fibers in 110 PubMed publications with DNA fibers published in 2023. **(d)** Examples of inter-user variability in the segmentation of disconnected or spotty fibers. Fibers were segmented independently by three users. **(e)** Examples of inter-user variability in the segmentation of disconnected or spotty fibers. After 4 months, three independent users were asked to segment the same set of DNA fibers as in **(d)**. **(f)** A schematic representation illustrating the unintentional omission of short DNA fibers in manual DNA fiber quantification. **(g)** Examples of inter-user variability in the detection of short DNA fibers in manual DNA fiber quantification. Fibers were segmented independently by three users.

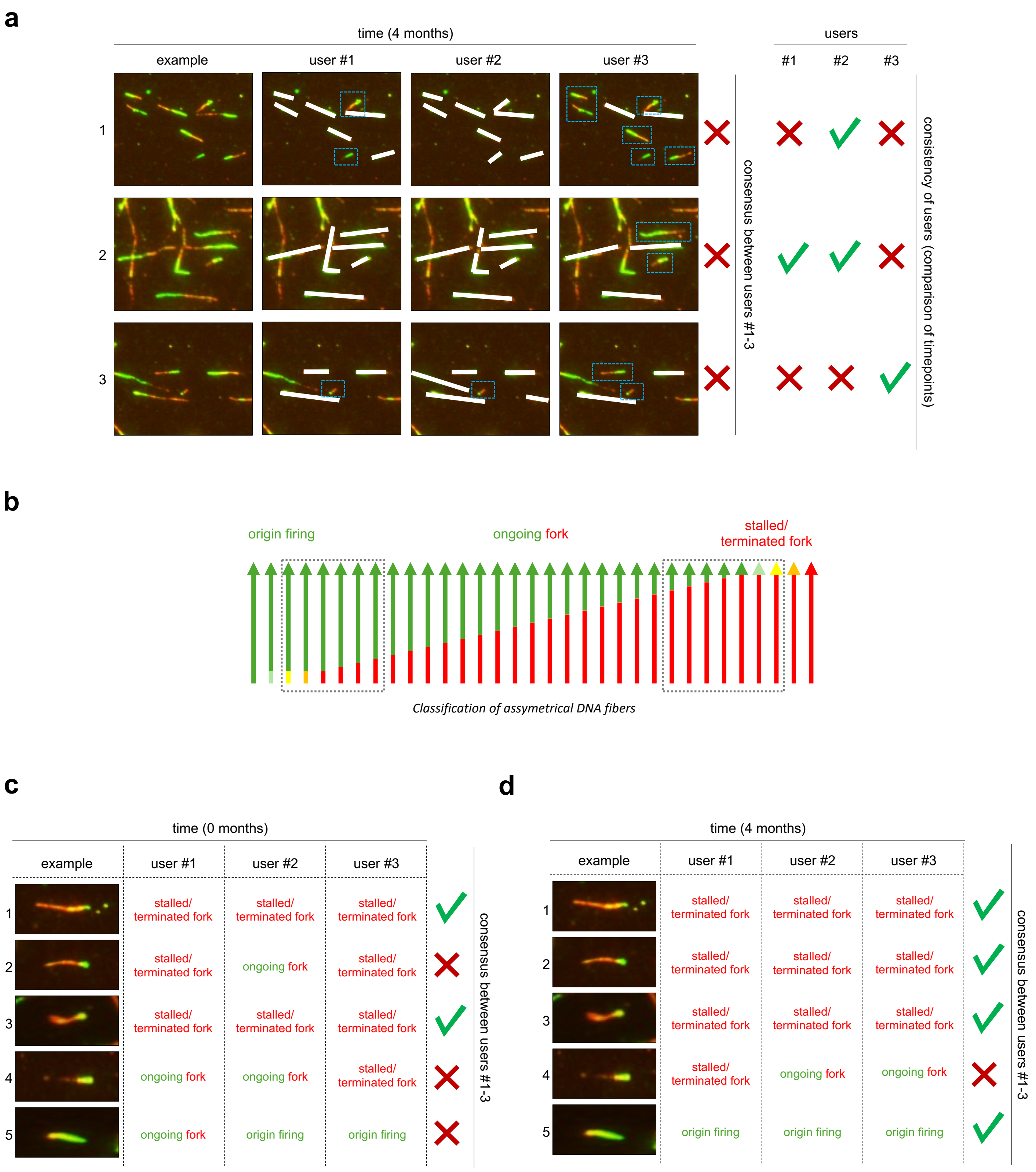


**Extended Data Figure 2**

**Extended Data Figure 2: Assessment of inter-user variability in the classification of asymmetrical DNA fibers.** **(a)** Examples of inter-user variability in the detection of short DNA fibers. After 4 months, three independent users were asked to segment the same set of DNA fibers as in Extended Data Fig. 1g. **(b)** A schematic representation illustrating the grey zones in the manual classification of asymmetrical DNA fibers. **(c)** Examples of inter-user variability in the manual classification of asymmetrical DNA fibers. Fibers were classified independently by three users. **(d)** Examples of inter-user variability in the manual classification of asymmetrical DNA fibers. After 4 months, three independent users were asked to classify the same set of DNA fibers as in **(c)**.

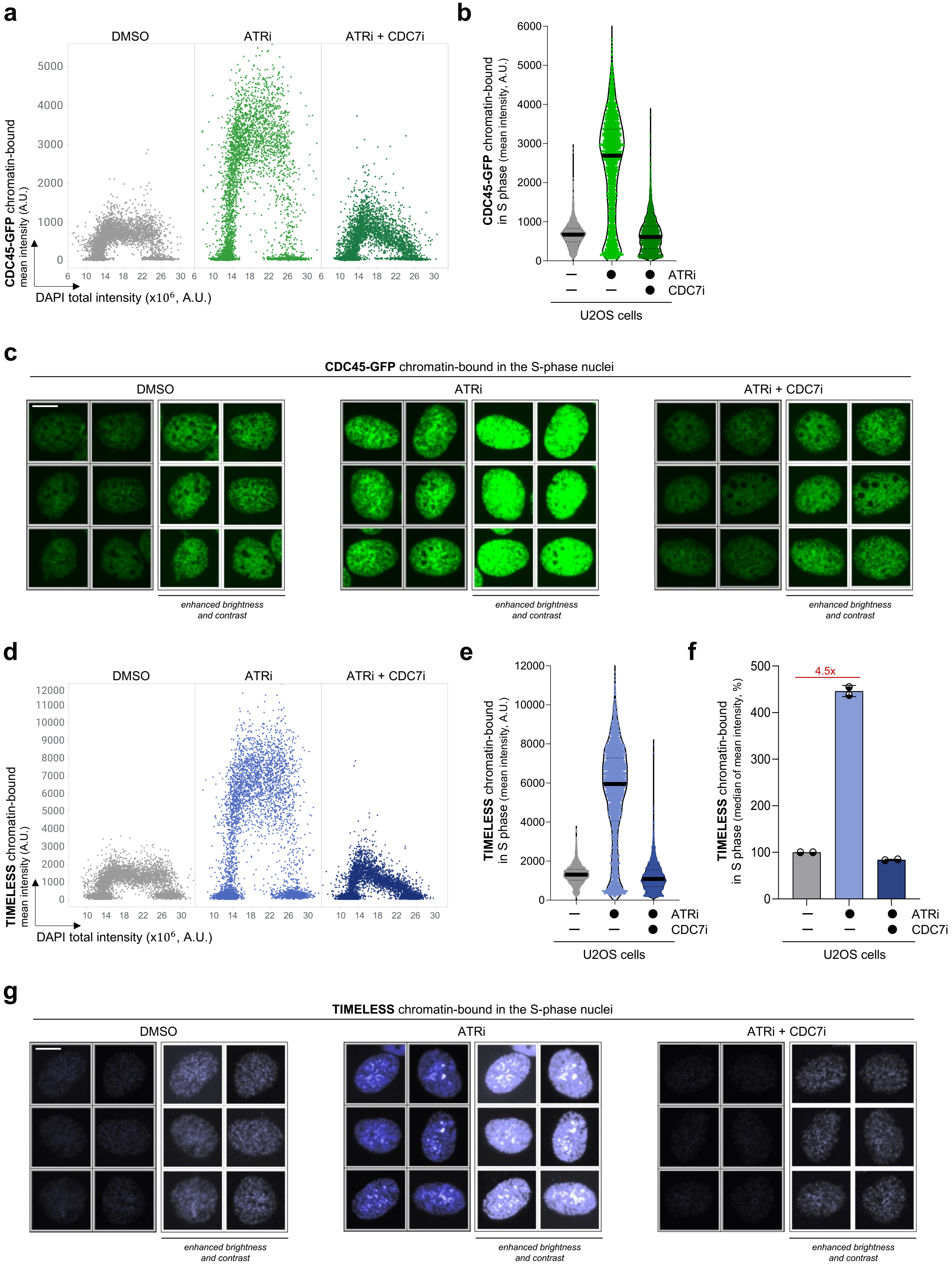


**Extended Data Figure 3**

**Extended Data Figure 3: QIBC analysis of CDC45 and TIMELESS, replisome components, in response to oscillations in origin firing. (a)** QIBC plots of pre-extracted U2OS cells stained for GFP after indicated treatment (5 µM ATRi, 5 µM CDC7i for 1 h). DAPI counterstains nuclear DNA. n ≈ 6,000 cells per condition. **(b)** Mean fluorescent intensity (MFI) of CDC45-GFP in S-phase cells based QIBC in **(a)**. Lines denote medians; n ≈ 3,000 cells per condition. **(c)** Unbiased QIBC galleries of CDC45-GFP U2OS cells immunostained for GFP upon indicated treatment. Scale bar, 10 µm. **(d)** QIBC plots of pre-extracted U2OS cells stained for TIMELESS after indicated treatment (5 µM ATRi, 5 µM CDC7i for 1 h). DAPI counterstains nuclear DNA. n ≈ 6,000 cells per condition. **(e)** MFI of TIMELESS in S-phase cells based QIBC in **(a)**. Lines denote medians; n ≈ 3,000 cells per condition. **(f)** Quantification of QIBC plots in **(d)**; each bar indicates the median of mean intensity (data are mean ± s.d.; n = 2 independent experiments). **(g)** Unbiased QIBC galleries of U2OS cells immunostained for TIMELESS upon indicated treatment. Scale bar, 10 µm. A.U. denotes Arbitrary Units.

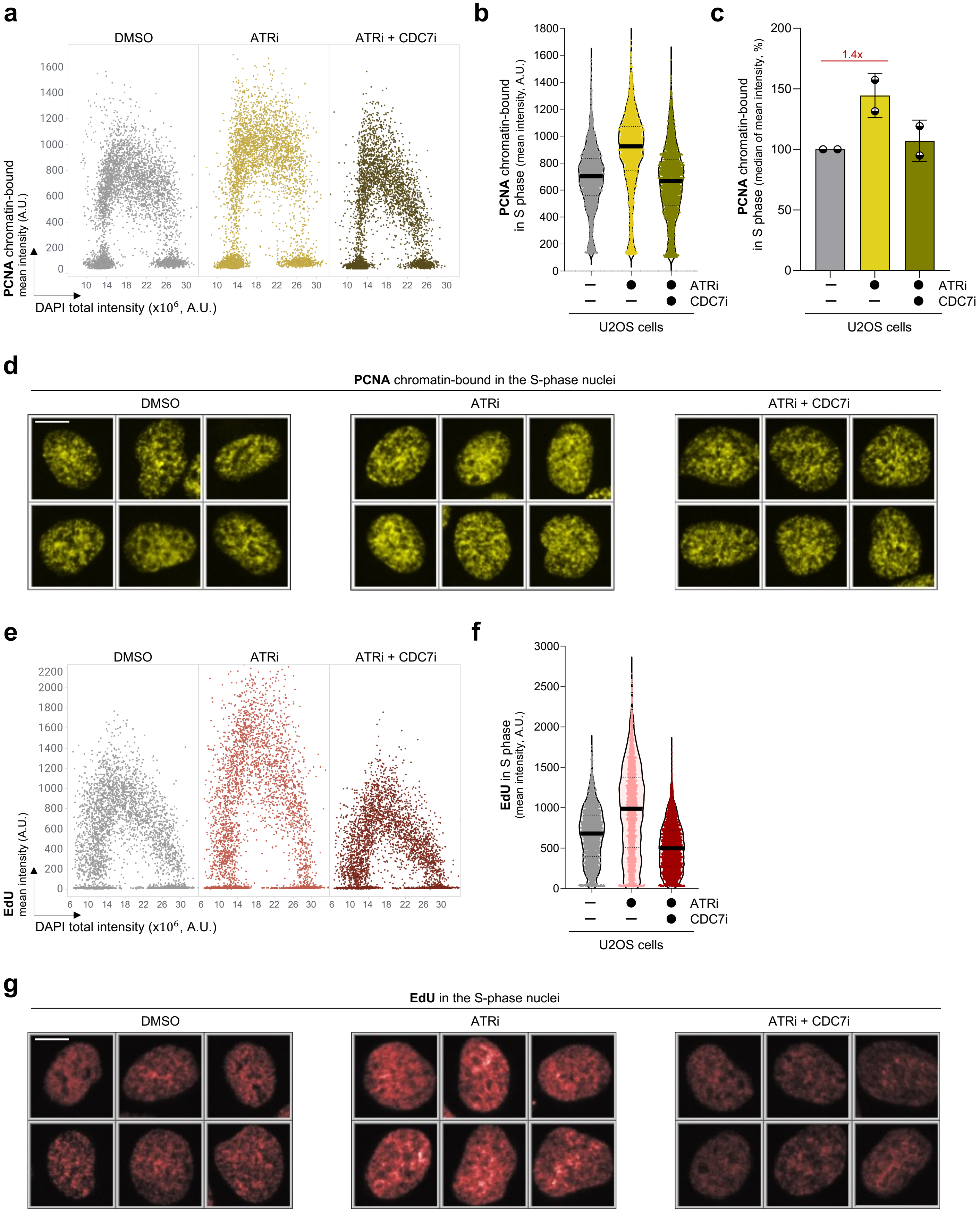
a
DMSO
ATRi
ATRi + CDC7i
PCNA chromatin-bound
mean intensity (A.U.)
DAPI total intensity (x10$^6$, A.U.)
b
PCNA chromatin-bound
in S phase (mean intensity, A.U.)
ATRi
CDC7i
U2OS cells
c
PCNA chromatin-bound
in S phase (median of mean intensity, %)
1.4x
ATRi
CDC7i
U2OS cells
d
PCNA chromatin-bound in the S-phase nuclei
DMSO
ATRi
ATRi + CDC7i
e
DMSO
ATRi
ATRi + CDC7i
EdU
mean intensity (A.U.)
DAPI total intensity (x10$^6$, A.U.)
f
EdU in S phase
(mean intensity, A.U.)
ATRi
CDC7i
U2OS cells
g
EdU in the S-phase nuclei
DMSO
ATRi
ATRi + CDC7i

**Extended Data Figure 4: QIBC analysis of PCNA component and DNA synthesis in response to oscillations in origin firing. (a)** QIBC plots of pre-extracted U2OS cells stained for PCNA after indicated treatment (5 µM ATRi, 5 µM CDC7i for 1 h). DAPI counterstains nuclear DNA. n ≈ 6,000 cells per condition. **(b)** MFI of PCNA in S-phase cells based QIBC in **(a)**. Lines denote medians; n ≈ 3,000 cells per condition. **(c)** Quantification of QIBC plots in **(a)**; each bar indicates the median of mean intensity (data are mean ± s.d.; n = 2 independent experiments). **(d)** Unbiased QIBC galleries of U2OS cells immunostained for PCNA upon indicated treatment. Scale bar, 10 µm. **(d)** QIBC plots of U2OS cells stained for EdU after indicated treatment (5 µM ATRi, 5 µM CDC7i for 1 h). DAPI counterstains nuclear DNA. n ≈ 6,000 cells per condition. **(e)** MFI of EdU in S-phase cells based QIBC in **(a)**. Lines denote medians; n ≈ 2,800 cells per condition. **(g)** Unbiased QIBC galleries of U2OS cells stained for EdU upon indicated treatment. Scale bar, 10 µm. A.U. denotes Arbitrary Units.

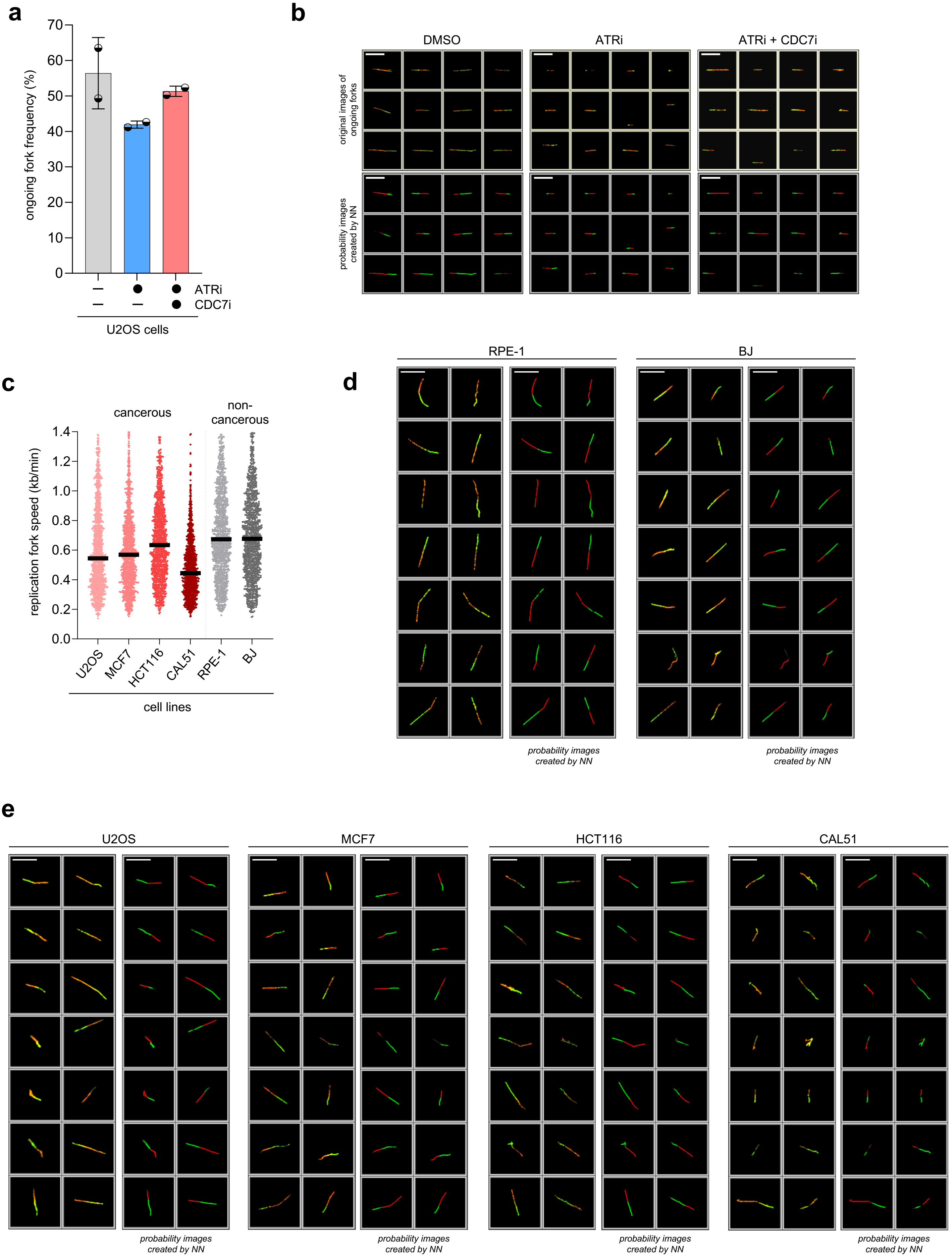


**Extended Data Figure 5**

**Extended Data Figure 5: qAID analysis of cell line panel containing non-cancerous and cancerous samples. (a)** qAID-based quantification of ongoing fork frequency in U2OS cells treated with indicated inhibitors (data are mean ± s.d.; n = 2 independent experiments). **(b)** Unbiased qAID galleries of DNA fibers in Fig. 2k. Both original fiber images and probability images created by the classification network are displayed. Galleries were masked based on segmentation. Scale bar, 10 µm. **(c)** qAID-based quantification of replication fork speed for ongoing forks in a panel of cancerous and non-cancerous cell lines. DNA fibers were prepared by spreading technique. Lines represent medians; n ≈ 1600 fibers per condition (except CAL51, n ≈ 500 fibers). **(d)** Unbiased qAID galleries of DNA fibers obtained from non-cancerous cells in **(c)**. Both original fiber images and probability images created by the classification network are displayed. Galleries were masked based on segmentation. Scale bar, 10 µm. **(e)** Unbiased qAID galleries of DNA fibers obtained from cancerous cells in **(c)**. Both original fiber images and probability images created by the classification network are displayed. Galleries were masked based on segmentation. Scale bar, 10 µm.

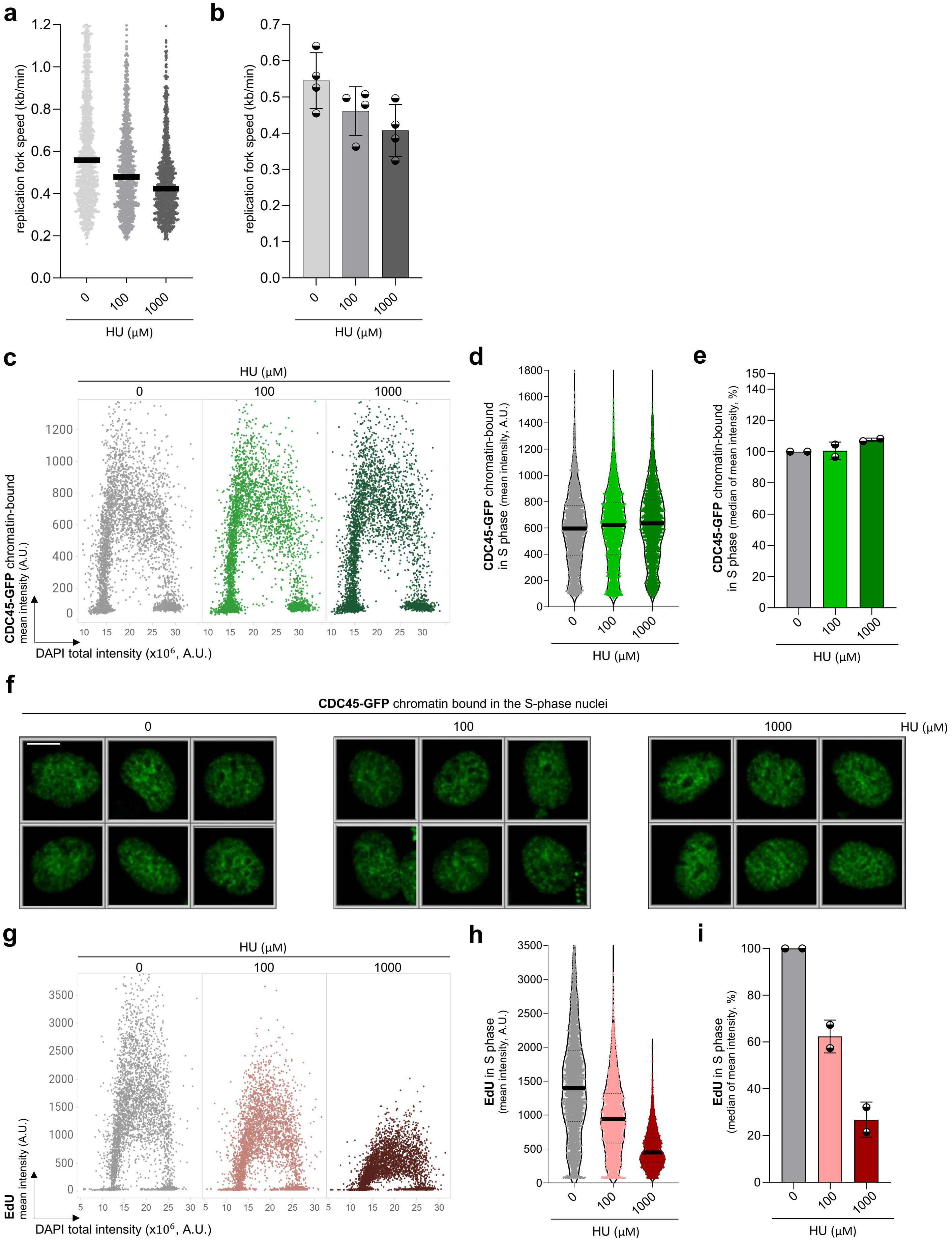


### Extended Data Figure 6

**Extended Data Figure 6: QIBC analysis of CDC45 and DNA synthesis in response to low and high HU concentrations. (a)** qAID-based quantification of replication fork speed for ongoing forks in U2OS cells with indicated HU treatment (1 h). DNA fibers were prepared by spreading technique. Lines represent medians; n ≈ 1000 fibers per condition. **(b)** Quantification of qAID-based replication fork speed plots in **(a)**; data are mean ± s.d.; n = 4 independent experiments. **(c)** QIBC plots of pre-extracted U2OS cells stained for GFP after indicated HU treatment (1 h). DAPI counterstains nuclear DNA. n ≈ 5,000 cells per condition. **(d)** MFI of CDC45-GFP in S-phase cells based QIBC in **(c)**. Lines denote medians; n ≈ 2,600 cells per condition. **(e)** Quantification of QIBC plots in **(c)**; each bar indicates the median of mean intensity (data are mean ± s.d.; n = 2 independent experiments). **(f)** Unbiased QIBC galleries of CDC45-GFP U2OS cells immunostained for GFP upon indicated treatment. Scale bar, 10 µm. **(g)** QIBC plots of U2OS cells stained for EdU after indicated HU treatment (1 h). DAPI counterstains nuclear DNA. n ≈ 5,000 cells per condition. **(h)** MFI of EdU in S-phase cells based QIBC in **(g)**. Lines denote medians; n ≈ 3,000 cells per condition. **(i)** Quantification of QIBC plots in **(g)**; each bar indicates the median of mean intensity (data are mean ± s.d.; n = 2 independent experiments). A.U. denotes Arbitrary Units.

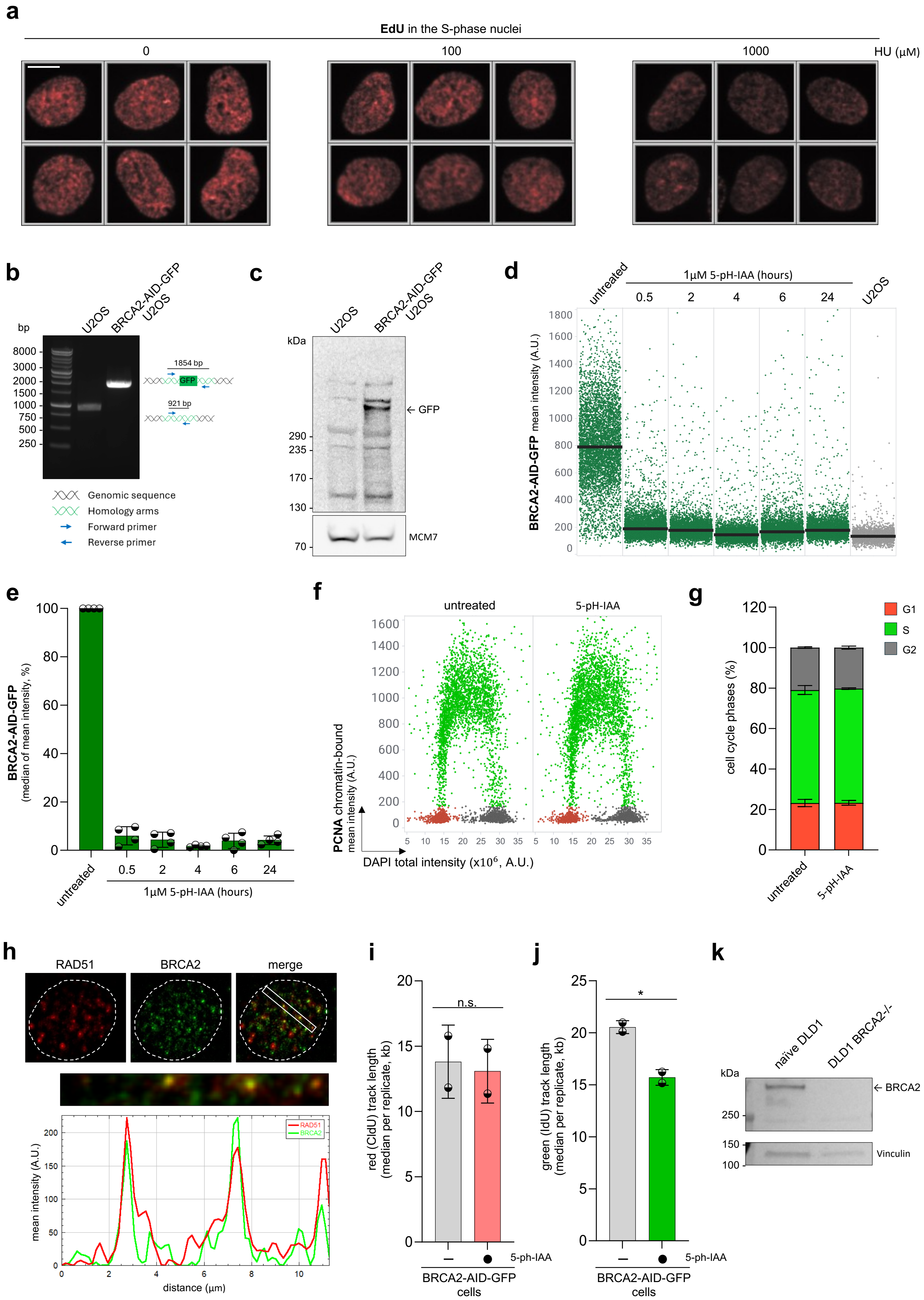


### Extended Data Figure 7

**Extended Data Figure 7: Generation and validation of BRCA2 degron in U2OS cells. (a)** Unbiased QIBC galleries of U2OS cells stained for EdU upon indicated HU treatment (1 h). Scale bar, 10 µm. **(b)** Junction PCR showing homozygous BRCA2-AID-GFP tagging. **(c)** Western blots of whole cell lysates obtained from naïve U2OS and BRCA2-AID-GFP U2OS cells stained for GFP. MCM7 was used as a loading control. **(d)** QIBC of naïve U2OS and BRCA2-AID-GFP U2OS cells stained for GFP after indicated 5-pH-IAA treatment. n ≈ 5,000 cells per condition. **(e)** Quantification of QIBC plots in **(d)**; each bar indicates the median of mean intensity (data are mean ± s.d.; n = 4 independent experiments). **(f)** QIBC plots of pre-extracted BRCA2-AID-GFP U2OS cells stained for PCNA after indicated 5-pH-IAA treatment. DAPI counterstains nuclear DNA; n ≈ 5,000 cells per condition. **(g)** Quantification of individual cell cycle phases based on QIBC in **(f)**; n = 2 independent experiments. **(h)** Representative widefield images of BRCA2 and RAD51 foci in U2OS cells supplemented by profile plot display the colocalization of BRCA2 and RAD51. **(i)** Quantification of qAID-based red track length plots in Fig. 3k; data are mean ± s.d.; n = 2 independent experiments. P values were calculated by a two-tailed unpaired *t*-test, not significant (n.s.) denotes P > 0.1. **(j)** Quantification of qAID-based green track length plots in Fig. 3k; data are mean ± s.d.; n = 2 independent experiments. P values were calculated by a two-tailed unpaired *t*-test, *P=0.0196, not significant (n.s.) denotes P > 0.1. **(k)** Western blots of whole cell lysates obtained from naïve and BRCA2-knockout DLD1 cells stained for BRCA2. Vinculin was used as a loading control. A.U. denotes Arbitrary Units.
